# Reusing pipettes in Patch-seq: a foundation for automated multimodal profiling

**DOI:** 10.64898/2026.11.10.763771

**Authors:** Luke Campagnola, Martin Chase, Jessica Trinh, Megan Koch, Audrey McCutcheon, Miki Lee, Brian Lee, Rachel Dalley, Rusty Mann, Alex Cahoon, Thomas Braun, Kathryn Gudsnuk, Faith Best, Staci Sorensen, Tim Jarsky

## Abstract

Patch-seq is a technique for single-cell multimodal characterization. It combines whole-cell electrophysiology, single-cell transcriptomics, and morphological reconstruction in single neurons. Patch-seq morphologies are a critical bridge among cell types, genes, physiology, and large-scale connectivity platforms such as EM connectomes and single-cell-resolution projectomes. However, patch-seq experimental complexity and operator dependence limit throughput. Automation could enable scalable Patch-seq but requires procedural adaptations, including detergent-based pipette cleaning, robotic nucleus extraction and collection. The impact of these automations on Patch-seq quality has not been systematically evaluated. Here, we show that each of these modifications preserves or improves key Patch-seq outcomes. Automated nucleus extraction performs comparably to, or better than, expert manual operation, preserving key RNA and morphological metrics. Detergent-cleaned, reused pipettes supported a viable sealing success rate across multiple cleaning cycles and preserved electrophysiological quality and transcriptomic yield without introducing excess contamination. Sonication effectively allows collection of nuclear material without damaging RNA and supports subsequent pipette cleaning and reuse. Enhanced automated sealing and break-in procedures increased throughput regardless of brain region, cell type or species. Together, these results validate the core procedural adaptations required for automated Patch-seq and establish a foundation for high-throughput, standardized multimodal single-cell characterization.

## Introduction

Upcoming large-scale descriptions of neuronal connectivity^1^ are poised to transform our understanding of brain circuitry in the same way that single-cell transcriptomics has transformed our understanding of cell-type diversity^2^. Dense synaptic reconstructions, including datasets such as MICrONS^3^, now provide an increasingly detailed view of local circuit structure, and projection datasets^4,5^ map the long-range axonal targets of defined neuronal populations. However, the value of these resources depends on our ability to relate connectional structure to defined cell types that underlie circuit function. Without such linkages, wiring diagrams remain difficult to interpret in terms of molecular programs, physiological properties, and function. Importantly, connectivity maps that lack molecular data are unable to inform circuit manipulation via molecular techniques, a strategy that has already demonstrated its utility in the Drosophila^6^.

Patch-seq is well positioned to provide this bridge by linking the gene expression profiles of single neurons to their intrinsic electrophysiological properties and local morphology, enabling multimodal cell characterization at the single-cell level^7^. These measurements are particularly valuable because local morphology provides features that can be aligned across experimental platforms. Thus, Patch-seq offers a path for relating transcriptomically defined cell types to morphologies recovered from electron microscopy datasets and, thereby, to synaptic connectivity^8^. The same logic applies to large projection datasets^9^. Although the long-range axons of projection neurons are truncated in slice-based Patch-seq experiments, the local morphologies captured by Patch-seq can still serve as a point of correspondence to full-morphology reconstructions and other projection-oriented datasets, helping link transcriptomic type and gene expression to long-range connectivity patterns.

The increasing importance of these cross-platform linkages has heightened the need for Patch-seq datasets at a much larger scale. However, Patch-seq remains technically challenging and relatively low-throughput, making it difficult to reach the scale required to support broad integration with extensive connectomic and projectomic datasets. Full automation of Patch-seq, by enabling parallelization, offers a path to overcoming the scalability limitations of manual experiments. Although automation promises clear advantages^10–15^, it also raises important concerns regarding compatibility with Patch-seq. Procedural adaptations required for full automation, such as detergent-based pipette cleaning^16,17^, sonication-based nucleus collection, automated nucleus extraction, and enhanced seal and break-in algorithms, risk compromising data quality and experimental performance. Because the value of Patch-seq depends on preserving the integrity of electrophysiological, molecular, and morphological measurements, it is necessary to determine whether these modifications alter the quality and quantity of the resulting data.

Here, we tested whether key adaptations required for automating Patch-seq are compatible with successful multimodal cell characterization. We investigated three primary areas: (1) whether automated nucleus extraction can match or exceed the success rates of expert operators, especially at preserving morphological quality; (2) whether sonication-based nucleus collection generates sufficient levels of high-quality RNA; and (3) whether detergent-based cleaning allows pipette reuse without excessive RNA cross-contamination between cells. We also evaluate whether changes to our seal and break-in algorithms improve throughput across various species, cell types, and brain regions.

## Results

To test the impact of our new patch-seq automation methods, we used our existing pipeline, in which several patch-clamp rigs operate in parallel on brain slices from a variety of mouse, human, and nonhuman primate (NHP) donors. We distributed slices from the same donor across our standard and automation-enhanced rigs to allow unbiased comparison between workflows. Our standard workflow (Fig. 1, orange arrow pathway), requires a new pipette for each patching attempt. After manually contacting the cell, sealing, break-in, and whole-cell electrophysiological recording are handled automatically with our standard algorithms. After enough time to allow biocytin to fill the cell, the nucleus is extracted manually and the pipette tip is broken off into a PCR tube to allow RNA sequencing. In contrast, the automation-enhanced workflow (Fig. 1, blue arrow pathway) uses updated algorithms for sealing and break-in and completely automates the process of extracting nuclei and depositing them into RNA collection tubes without breaking the pipette tip, allowing the pipette to be cleaned and reused for subsequent patching attempts.

**Figure 1.**
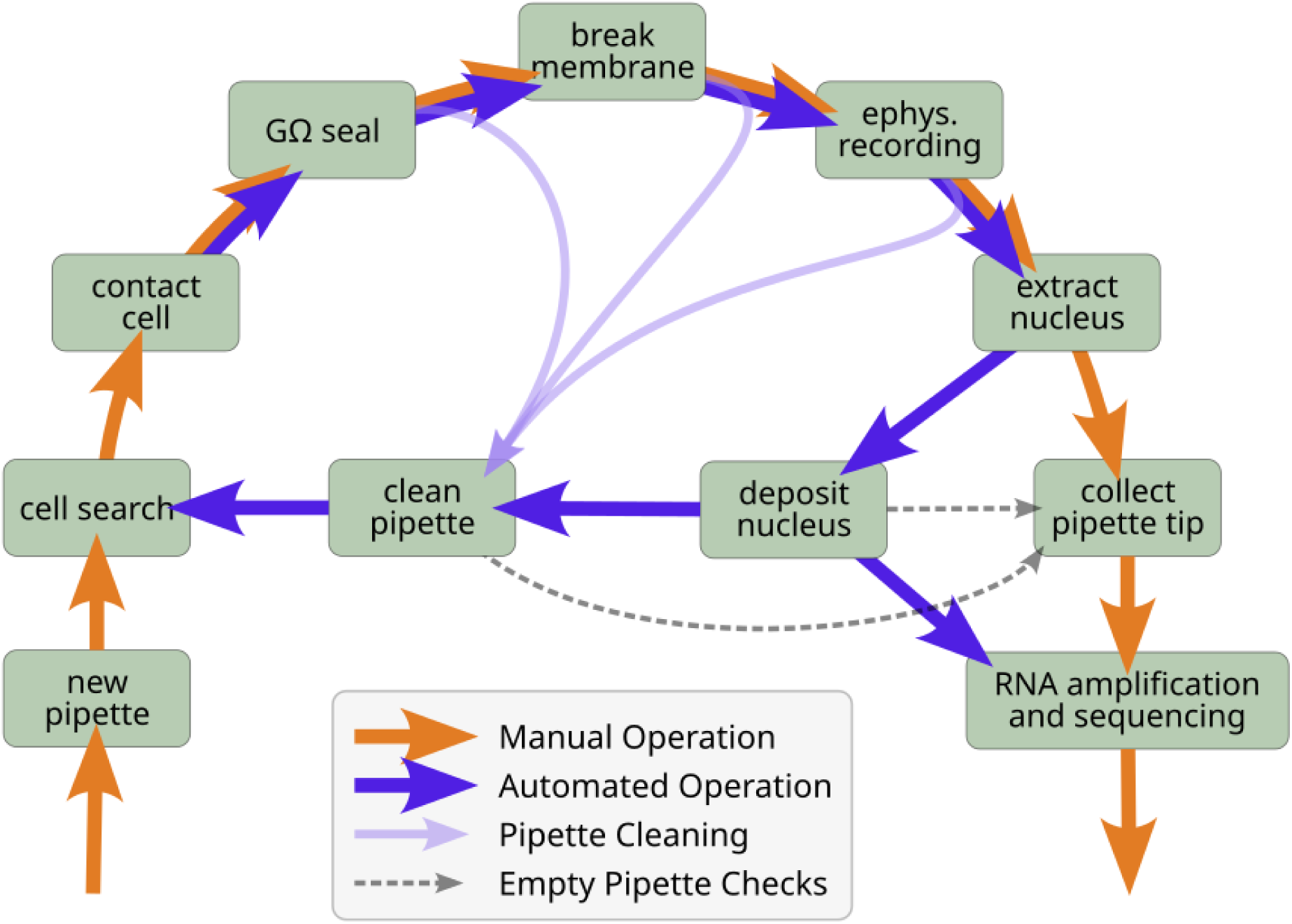
Manual and automatic Patch-seq workflow. Flowchart shows key stages in the process of a single attempted Patch-seq recording. Orange arrows indicate the manual workflow whereas blue arrows indicate automated workflow. Light blue arrows indicate possible patch-failure paths where a pipette would be discarded in the manual workflow, but can instead be reused when pipette cleaning is available. Dashed grey arrows indicate negative control experiments where a pipette is tested for residual RNA after being cleaned, or directly after depositing a collected nucleus.

We first compared standard-versus enhanced-workflow performance during gigaohm (GΩ) seal formation and whole-cell break-in (see Methods). Although the enhancement of these steps was not required to achieve Patch-seq automation, it provided a low-risk entry point for us to establish controlled testing of updates to the workflow. To ensure matched comparisons, seal analyses were restricted to donors (mouse, human, or nonhuman primate) with at least one automated seal attempt, and break-in analyses were restricted to successfully sealed attempts from donors with at least one automated break-in attempt.

Our enhanced automated sealing procedure increased the proportion of successful GΩ seal attempts (Fig. 2A). Standard operation succeeded in 76% of attempts (1043/1371), whereas enhanced operation produced GΩ seals in 88% (738/839; fisher exact p=2e-12). Enhanced break-in also improved success rates (Fig. 2B). Among successful seals, standard break-in yielded a healthy whole-cell configuration in 82% of attempts (907/1100), whereas enhanced automated break-in succeeded in 90% (466/516, fisher exact p=3e-5). Because these stages occur in sequence, the improvements compound, raising the estimated whole-cell yield per sealing attempt from 62% to 79%.

**Figure 2.**
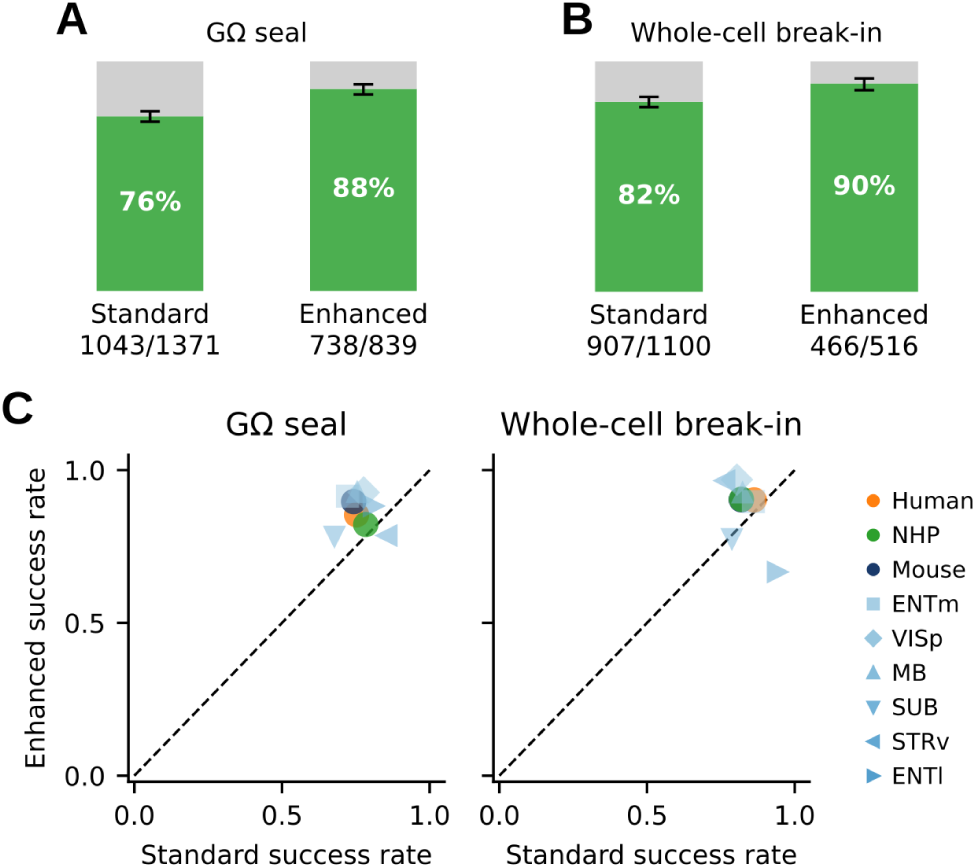
Automated sealing and break-in improve performance over standard method. **A.** Proportion of patch attempts achieving a GΩ seal for manual and automated approaches. Only cells with at least one automated seal attempt are included. **B.** Proportion of GΩ sealed attempts with successful break-in to whole-cell configuration. Cells without automated break-in attempts are excluded. Error bars in (A) and (B) indicate 95% Wilson confidence intervals. **C.** Per-group seal (left) and break-in (right) success rates comparing standard (x-axis) to enhanced (y-axis) approaches. Circles represent species (Human: orange; NHP: green; Mouse: dark blue). Other shapes represent mouse brain regions represented in the data (shades of lighter blue). Dashed diagonal indicates equal performance.

Furthermore, performance in these key steps was unaffected by either the species or brain region of the sample (Fig. 2C).

Next, we assessed whether detergent-based pipette cleaning, which enables pipette reuse during automated workflows, affects sealing success, electrophysiological measurements, RNA recovery, and transcriptomic mapping.

To test whether detergent cleaning could support pipette reuse without compromising Patch-seq outcomes, we compared recordings obtained with fresh pipettes to those obtained with pipettes that had undergone one or more prior cleaning cycles. Analyses were restricted to donors that contributed at least one automated break-in attempt.

Fresh pipettes produced GΩ seals in 81% of attempts (591/731; Fig. 3A). After one prior cleaning cycle, this rate dropped to 59% (164/276), and after two or more prior cleaning cycles, 52% (24/46). These results show that detergent-cleaned pipettes remain usable across multiple cleaning cycles, although sealing success was reduced compared to fresh pipettes.

**Figure 3.**
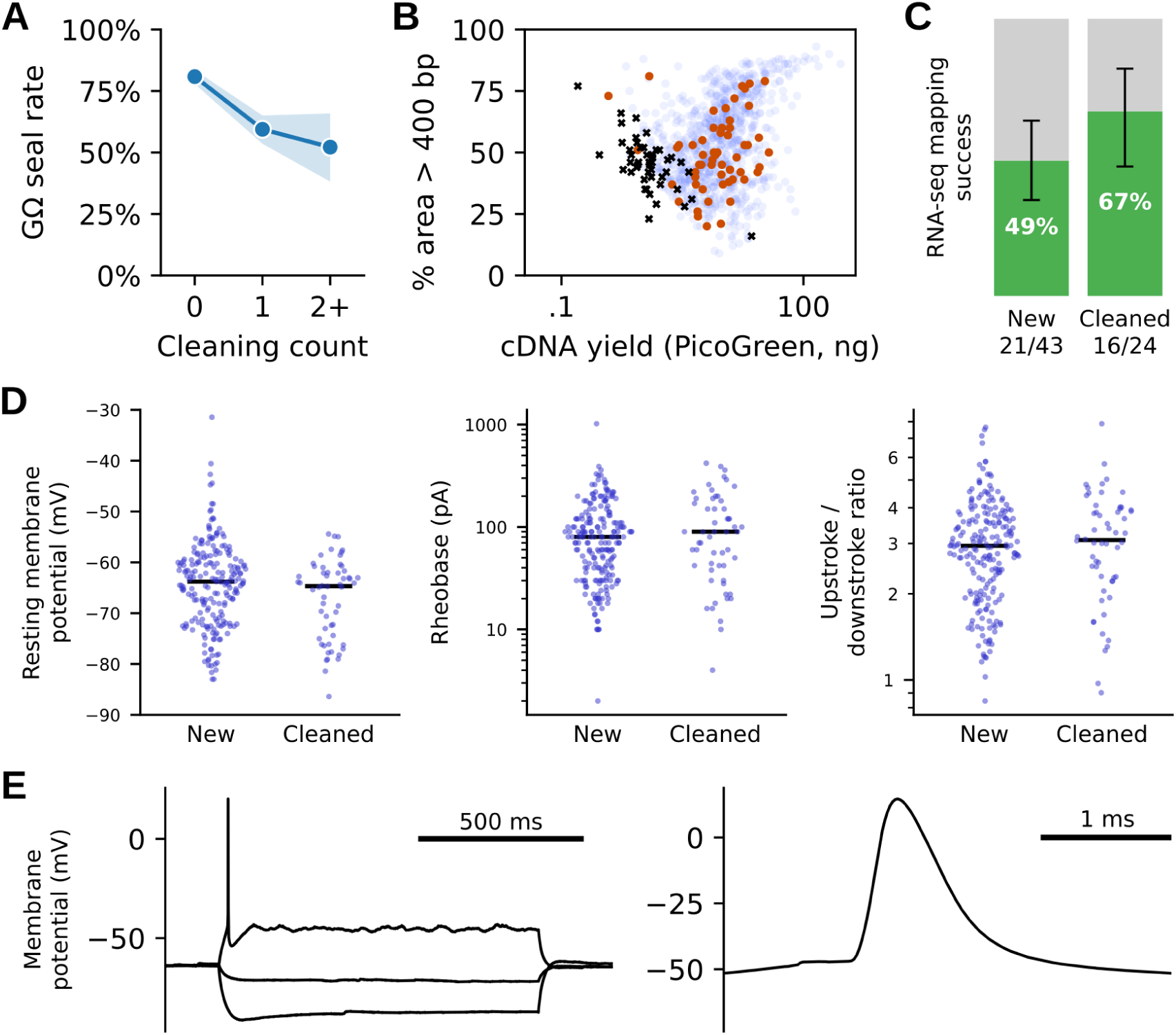
Pipette cleaning with detergent is compatible with Patch-seq. All panels include only donors for which at least one patch attempt was performed with a previously-cleaned pipette. **A.** Proportion of patch attempts achieving a GΩ seal, grouped by the number of times the pipette had been cleaned before the attempt (0 = new, 1, or ≥2 prior cleans). Filled region indicates 95% Wilson score confidence intervals. **B.** cDNA quality (percentage of amplified fragments > 400 bp) versus cDNA yield (total amplified cDNA measured by PicoGreen, pg; log scale) for cells collected with new pipettes (blue) versus previously-cleaned pipettes (red). Negative controls (black X) generated by sequencing from pipette tips that had been cleaned after patching a cell but failing to extract a nucleus. Attempts using automatic nucleus deposit are excluded, as are samples whose Fragment Analyzer trace showed a capillary step artifact. **C.** Proportion of sequenced nuclei with high-quality cell type mapping. Attempts using automatic nucleus deposit are excluded. **D.** Intrinsic electrophysiological properties of cells recorded with fresh versus previously-cleaned pipettes: resting membrane potential (mV, linear scale), rheobase (minimum current to elicit a spike, pA, log scale), and action potential upstroke/downstroke ratio (log scale). Lines show group medians; points show individual cells. **E.** Representative recordings from a mouse CA3 neuron made with a cleaned and reused pipette.

To assess compatibility with Patch-seq molecular recovery, we compared cDNA yield and Fragment Analyzer-based cDNA quality metrics for samples collected with fresh versus cleaned pipettes, excluding samples collected via automatic nucleus deposit. cDNA yield, measured by PicoGreen, and cDNA quality, assessed as the percentage of amplified fragments greater than 400 bp, were similar between groups (Fig. 3B). We also observed that the ability to map mRNA from Patch-seq samples to single-cell-transcriptomics based taxonomies^18^ was preserved (Fig. 3C). Thus, detergent cleaning produced no evidence of reduced or degraded RNA recovery among successful recordings.

We next examined whether use of a cleaned pipette altered basic intrinsic electrophysiological measurements. Resting membrane potential, rheobase, and action potential upstroke/downstroke ratio were broadly similar for cells recorded with fresh versus cleaned pipettes (Fig. 3D & E), indicating that detergent cleaning did not produce obvious distortion of the intrinsic physiological properties examined here.

Together, these results indicate that detergent-based pipette cleaning is compatible with Patch-seq. Cleaned pipettes can be reused for successful recordings and preserve both electrophysiological measurements and downstream cDNA quality, supporting their use in automated workflows that reduce interruptions from manual pipette replacement.

We next evaluated whether automated nucleus extraction, among the most technically demanding and critical stages of Patch-seq, could be implemented without reducing sample throughput, morphology recovery, or RNA recovery. During this stage, the pipette is withdrawn while suction is applied to hold the nucleus. Careful retraction results in the cell body remaining in place while the membrane pinches off and re-seals around the nucleus, thereby preventing loss of RNA that is critical for mapping to existing taxonomies^19^. Furthermore, successfully resealing the cell membrane during retraction prevents leakage of biocytin, preserving the ability to stain the cell for morphological reconstruction. Analyses of reseal success were restricted to donors with at least one automated extraction attempt and to patch attempts that achieved a successful whole-cell recording.

As an initial measure, experimenters checked visually for the presence of a nucleus at the pipette tip after manual and automated extraction (Fig. 4A). Manual extraction successfully recovered a nucleus in 50% of attempts (664/1337), whereas automated extraction improved nucleus recovery, succeeding in 59% of attempts (396/673; fisher exact p=1e-4).

**Figure 4.**
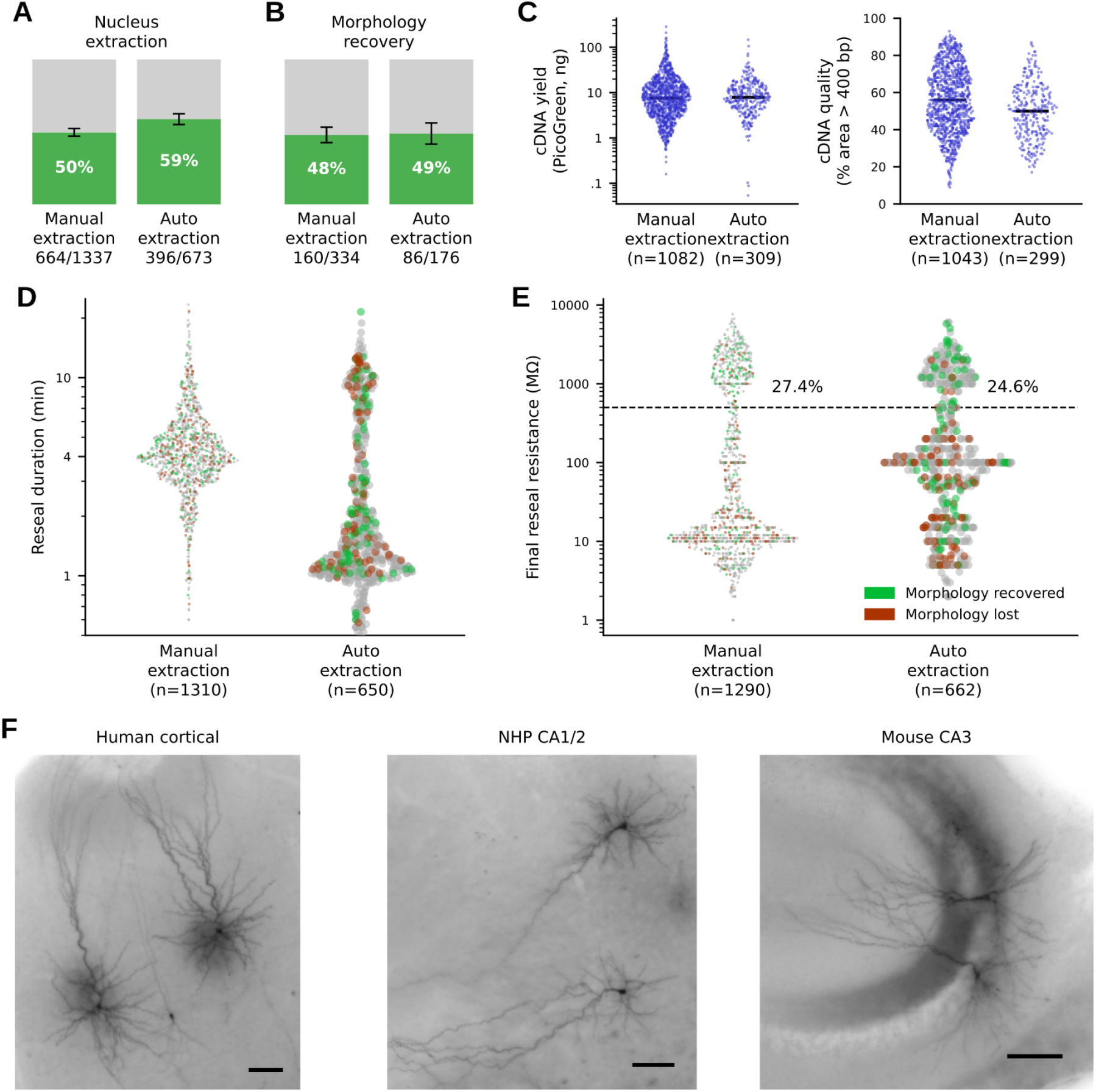
Automated nucleus extraction performs as well as or better than manual extraction. **A.** Proportion of nucleus extraction attempts where a nucleus was observed at the pipette tip after retraction for manual (left) and automated (right) extraction. Error bars are 95% Wilson binomial confidence intervals. **B.** Proportion of attempts that yielded high quality cell morphology. **C.** cDNA yield (total amplified cDNA measured by PicoGreen, pg; log scale) and quality (percentage of amplified fragments > 400 bp, linear scale) for manual and automated extraction, excluding attempts that used automatic nucleus deposit or where the Fragment Analyzer trace showed a step artifact. **D.** Reseal duration (log scale) for manual and automated extraction. Each point is one patch attempt, colored by morphology recovery. **E.** Final pipette resistance measured after retraction (log scale), with points colored as in (D). The dashed line marks a 500 MΩ reseal threshold; percentages indicate the fraction of attempts in each group meeting or exceeding this threshold. **F.** Representative images of biocytin-stained cells that survived automatic nucleus extraction.

Encouraged by this result, we next examined whether cellular morphology was preserved after resealing by assessing the proportion of extracted cells that yielded a high-quality morphology. After fixation and biocytin staining, quality was assessed using minimum intensity projection images acquired with a 10X or 20X objective. This assessment served as a gate for subsequent high resolution imaging with a 63X objective (Fig. 4F). High-quality morphologies were obtained for 48% of manually extracted cells (160/334) compared to 49% of automatically extracted cells (86/176; Fig. 4B). Thus, although automated extraction increased the rate of nucleus recovery, the rate of morphological recovery was not significantly different.

To assess compatibility with Patch-seq molecular recovery, we compared cDNA yield and Fragment Analyzer-based cDNA quality metrics for samples collected with manual versus automated extraction, excluding samples collected via automatic nucleus deposit. cDNA yield (PicoGreen, pg), was similar between groups (Fig. 4C; manual=7.5 ng, automated=7.8 ng; Mann-Whitney U p=0.3), whereas cDNA quality, assessed as the percentage of amplified fragments greater than 400 bp, was significantly lower in the automated group (manual=56%, automated=50%, Mann-Whitney U p=2e-7); however, the distributions largely overlap and the difference may have little practical effect.

To further evaluate the automatic nucleus extraction method, we compared both the duration of extraction and the final reseal resistance between manual and automated attempts. Both the duration of the resealing phase (Fig. 4D) and the final reseal resistance (Fig. 4E) showed overlapping distributions. Mean extraction time was lower for automated extraction (manual=4 minutes, automated=1.4 minutes; Mann-Whitney U p<1e-10) and final reseal resistance was higher (manual=20 MΩ, automated=100 MΩ; Mann-Whitney U p=6e-10), suggesting improved performance in both metrics by the automated approach. However, the final fraction of cells that had a final reseal resistance > 500 MΩ was not significantly different between the two (manual=25%, automated=27%; Fisher exact p=0.21) and notably, the bulk of successfully recovered morphologies are in these high-reseal resistance groups. Overall these results show comparable performance between manual and automated approaches, but hint that further optimization is possible. Together, these results show that automated nucleus extraction produces similar results to the standard manual approach, and is a viable step towards Patch-seq automation.

Finally, we assessed whether extracted nuclei can be automatically collected into a PCR tube without breaking the pipette tip. We found that high pressure and the properties of the lysis buffer together are not sufficient to expel nuclei from the pipette tip; in most cases the tip would become clogged or significant nuclear material would remain attached to the pipette. To solve this problem, we sonicated pipette tips while in the PCR tube lysis buffer. In combination with high pressure, this method enabled us to reliably remove and collect most material from the pipette. Standard and automated collections had median PicoGreen yields of 6.9 ng and 6.8 ng, respectively, as compared to a median of 1.2 ng for control pipettes that had failed nucleus collection and were cleaned before tip breaking (Fig. 5A).

**Figure 5.**
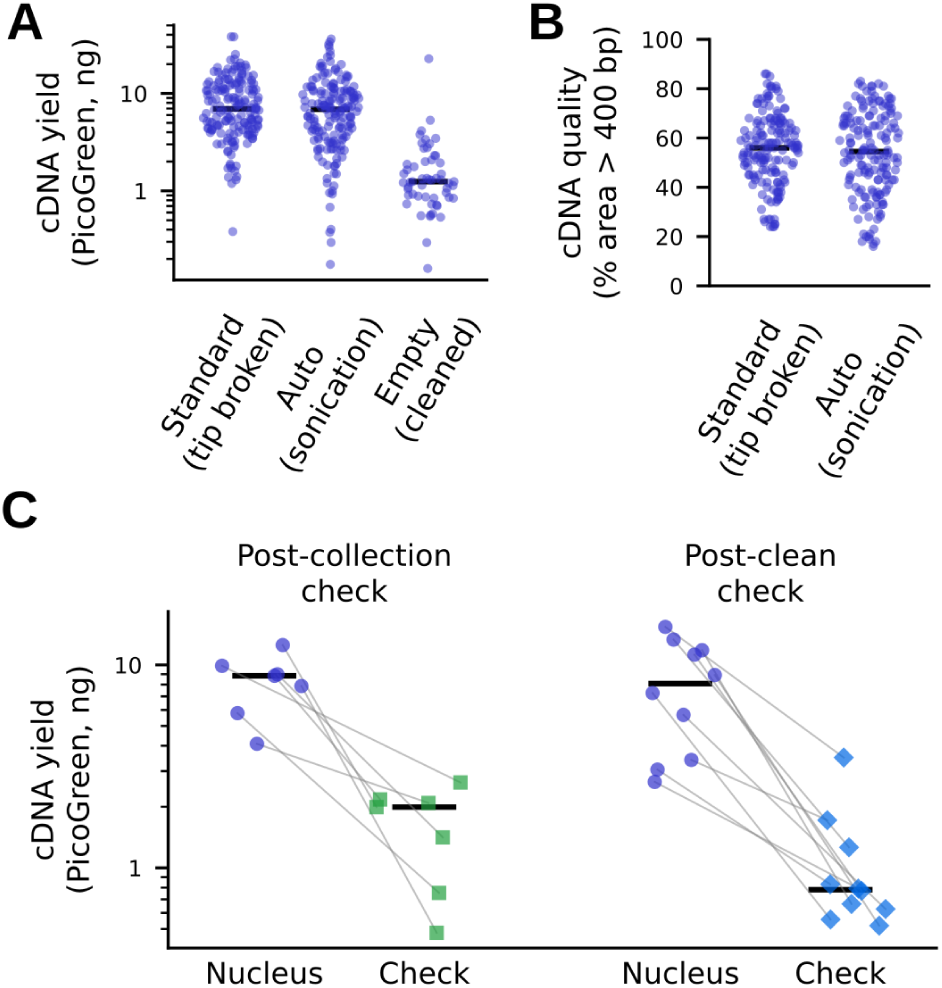
Pipettes can be reused after nucleus collection. All panels include only donors for which at least one patch attempt used automated nucleus deposition, and are restricted to attempts from which a nucleus was successfully extracted from the tissue. **A**. cDNA yield (total amplified cDNA measured by PicoGreen, pg; log scale) for nuclei deposited by the standard manual method (pipette tip broken into the collection tube) versus automated deposition using sonication and “empty” controls (pipettes that failed to collect a nucleus and were cleaned before breaking tip into the PCR tube). Bars show group median values and points are individual tubes. **B.** cDNA quality (percentage of amplified fragments > 400 bp) for the same 3 groups as in (A). **C**. cDNA quantity for automatically deposited nuclei (gray), compared with cDNA left in the same pipette immediately after nucleus collection (post-collection check, green) or after subsequent detergent cleaning (post-clean check, blue). Bars show group means; points show individual samples.

A natural concern with this approach is that ultrasound may directly shear RNA molecules, leading to degraded product. However, we found no significant difference in cDNA quality, measured as the fraction of amplified fragments in the 400 to 6,000 bp range, between the standard tip-breaking collection method and the automated sonication method. These results indicate that sonication-assisted automatic nucleus deposit can recover amplifiable RNA after nucleus extraction, supporting its use as an alternative to tip breaking in workflows designed to enable pipette reuse.

Our final question was whether pipette reuse would lead to RNA contamination in subsequent samples. To assess this, we collected nuclei using the automated method, followed by control measurements to determine the amount of residual RNA remaining in the pipette. In one group of controls, the pipette tip was broken into a second PCR tube immediately following nucleus collection (Fig. 5C; ’post-collection check’). Control tubes in this group contained significantly less cDNA (median 1.9 ng versus 8.8 ng; Wilcoxon signed-rank p=0.016), demonstrating that we are able to collect most RNA from the pipette, leaving very little behind.

To estimate the amount of RNA that would potentially contaminate the next sampled cell, we measured a similar group of controls, but following pipette cleaning (Fig 5C; ’post-clean check’). Cleaned pipettes contained near baseline levels of cDNA (median 0.8 ng versus 8.1 ng; Wilcoxon signed-rank p=0.002), showing that a pipette that is reused following a prior nucleus collection will add little extra contamination. Together, these results demonstrate a feasible path towards fully automated Patch-seq experiments.

## Discussion

Automation of patch clamp (and more recently Patch-seq) has been a longstanding goal in our field. Automation can increase throughput, reduce operator burden, and improve standardization^10,14^. However, the primary purpose of this project has been to improve *scalability*. In the past, patch clamp was fundamentally limited as a technique by its requirement to have near continuous human monitoring over a small number of carefully handled cells. A major potential of patch clamp automation, then, lies in the opportunity to dramatically scale into parallel data production once direct experimenter monitoring is no longer required.

Although patch clamp automation has made enormous strides in recent years, Patch-seq is particularly sensitive to the technical details of the method due to the requirement that it simultaneously maintain the quality of electrophysiological recording, morphological recovery, and RNA sampling. We have found that key procedural adaptations for automating Patch-seq not only match the quality and performance of manual operation, but in many cases lead to better metrics. Detergent-based cleaning allowed pipette reuse without affecting RNA, morphology, or electrophysiological features. Automatic nucleus extraction largely replicates the results of manual extraction. Nucleus expulsion assisted by sonication produced nuclear content similar to conventional tip-breaking without RNA damage. Finally, properly cleaned pipettes carry sufficiently low contamination to allow repeated collection of nuclei via the same reused pipette.

Pipette reuse^16,20^ is especially important because replacing pipettes requires either frequent human intervention or a complex mechanical solution; both are a major impediment to achieving high throughput, fully autonomous Patch-seq. Whereas detergent cleaning has previously been shown to have no significant effects on electrophysiological properties^16,17^, it was not obvious to us that trace amounts of detergent would not cause either RNA degradation or membrane degradation leading to decreased morphological recovery. Likewise, it was not clear that sufficient RNA capture would be possible without breaking the pipette tip for collection (indeed, the possibility of automating Patch-seq as been explicitly passed over previously for this reason^21^). Our results show that sonication-assisted nucleus collection combined with detergent-based cleaning can be used in Patch-seq without causing transcriptomic damage or excess carryover contamination. These findings support pipette cleaning as both a practical way to increase throughput and a method compatible with Patch-seq’s molecular requirements.

The goal of automation here is to enable scalability, which is important because Patch-seq’s role is expanding. It is no longer just a single-cell method for local cell-type characterization, but increasingly serves as a bridge linking cell morphologies seen in large datasets to their electrophysiological and transcriptomic profiles^8,9^. Large-scale light- and electron-microscopic reconstructions now reveal the projection patterns and synaptic connectivity of thousands to millions of neurons at a time, but only as static anatomical snapshots. Using cell morphology as a mapping, Patch-seq data allows us to predict the gene expression and intrinsic physiological properties of individual neurons in these massive datasets, and to link them to a much broader field of literature that operates on transcriptonic cell types. Improving Patch-seq scalability is therefore not just a technical advance in electrophysiology, but a necessary step for establishing physiological interpretation of large connectomics datasets.

The next step is to combine these validated components into more parallelized workflows, where one operator manages multiple pipettes or experimental stages at once. For automation to have a wide impact, the methods, software, and hardware must be accessible outside specialized labs. Automated Patch-seq will be most significant when paired with full-neuron morphology pipelines and computational tools for cross-platform integration, enabling the multimodal single-cell datasets needed to relate transcriptomic identity to morphology and connectivity at scale.

## Methods

### Patch-seq experiments

The Patch-seq workflow used in this study follows the previously established Allen Institute Patch-seq pipeline^19^, except where modifications were made to test automation. For this study, we performed Patch-seq using our standardized pipeline, while modifying a subset (2 of 8) of rigs to incorporate automation. The addition of various automation steps was progressive, allowing us to test one adaptation at a time. This comparative setup enabled us to assess the performance of the existing pipeline against that of the automation-enhanced rigs, using tissue from the same animals or surgical specimens.

### Mouse husbandry

All procedures were carried out in accordance with the Institutional Animal Care and Use Committee at the Allen Institute for Brain Science. Animals (<5 mice per cage) were provided food and water ad libitum and maintained on a regular 12-hr light–dark cycle. Animals were maintained on the C57BL/6 J background, and newly received or generated transgenic lines were backcrossed to C57BL/6 J.

### Non-human primate

We obtained non-human primate brain specimens, Macaca Nemestrina, Macaca Mulatta, and Macaca fascicularis, from animals designated for the Washington National Primate Research Center’s Tissue Distribution Program. Monkeys were housed in individual cages on a 12h light/dark cycle in a temperature and humidity-controlled room. All procedures involving non-human primates were approved by the University of Washington’s Institutional Care and Use Committee (IACUC) and conformed to the NIH’s Guide for the Care and Use of Laboratory Animals.

### Human

Human neurosurgical specimens were obtained from local hospitals in collaboration with neurosurgeons. Data included in this study were obtained from neurosurgical tissue resections for the treatment of refractory temporal lobe epilepsy or deep-brain tumors. All patients provided informed consent, and experimental procedures were approved by the hospital’s institutional review boards before the study commenced.

### Brain slice preparation

For mouse experiments, male and female mice were used between P45 and P70, anesthetized with 5% isoflurane, and intracardially perfused with 25 mL of 0–4°C slicing ACSF. Human, mouse, or macaque slices (350 μm) were generated (Compresstome VF-300 vibrating microtome, Precisionary Instruments or VT1200S Vibratome, Leica Biosystems), with a block-face image acquired (Mako G125B PoE camera with custom integrated software) before each section to aid in registration to the common reference atlases. Slices were transferred to an oxygenated and warmed (34 °C) slicing ACSF for 10 min, then transferred to room temperature holding ACSF (continuously bubbled with 95 % O2/5 % CO2) for the remainder of the day until transferred for patch-clamp recordings.

### Slice recording conditions

Before recording, all surfaces, equipment, and materials were sequentially cleaned with DNA Away (Thermo Scientific), RNase Zap (Sigma-Aldrich), and nuclease-free water. Acute brain slices (350 μm) were allowed to recover for at least 45 min at room temperature in the incubation chamber before recording. During recording, slices were bathed in warmed (34 °C) ACSF continuously bubbled with 95% O2/5% CO2. Recording ACSF contained 126 mM NaCl, 26 mM NaHCO3, 12.5 mM D-glucose, 2.5 mM KCl, 2 mM CaCl2·2H2O, 1 mM MgSO4, and 1.25 mM NaH2PO4·H2O, pH 7.3. To block fast synaptic transmission, the bath solution also contained 1 mM kynurenic acid and 0.1 mM picrotoxin.

### Cell targeting and recording hardware

Cells were visually targeted using oblique illumination contrast optics, in some cases in combination with fluorescence. Recordings were obtained using a Multiclamp 700B amplifier headstage (Molecular Devices) mounted on a PatchStar micromanipulator (Scientifica). Signals were acquired with an ITC-18 Data Acquisition Interface (HEKA). Command generation, signal processing, and amplifier metadata acquisition were performed in MIES (https://github.com/AllenInstitute/MIES), which runs in Igor Pro (WaveMetrics). Signals were low-pass filtered at 10 kHz with a Bessel filter and digitized at 50 kHz. Reported membrane potentials were not corrected for the measured liquid junction potential of −14 mV between the pipette and bath solutions.

### Pipette fabrication and internal solution

Recording pipettes were fabricated from thick-walled borosilicate glass (Warner Instruments, G150F-3) using a Narishige PC-10 puller and had a final resistance of 4-5 MΩ. Before recording, pipettes were filled with approximately 1.0-1.5 μL of internal solution containing 110 mM potassium gluconate, 10 mM HEPES, 0.2 mM EGTA, 4 mM KCl, 0.3 mM GTP sodium salt, 10 mM phosphocreatine disodium salt, 1 mM ATP magnesium salt, 20 μg/mL glycogen, 0.5 U RNase inhibitor (Takara, 2313A), and 0.5% biocytin (Sigma, B4261), adjusted to pH 7.3. Internal was filtered immediately before use to prevent pipette clogging during nucleus collection and pipette cleaning.

### Whole-cell recording procedure

Seal test monitoring and break-in were performed under voltage clamp. Whole-cell recordings were then carried out in current clamp. After formation of a stable seal and break-in, the resting membrane potential was measured, typically within the first minute. For the remainder of the recording, bias current was injected automatically during the interstimulus period using MIES algorithms to maintain the cell at its initial resting membrane potential. Bridge balance and pipette capacitance neutralization were adjusted manually in current-clamp. Series resistance compensation was not used. Stimulus paradigms were collected as previously described^19,22^.

### cDNA processing and analysis

For this study, RNA recovery was evaluated primarily using total cDNA yield and electropherogram trace analyses. Comparisons of detergent cleaning, sonication-based collection, and automated versus manual procedures were therefore based primarily on the quantity and quality of recovered and amplified RNA, rather than on downstream transcriptomic mapping or cell-type assignment.

For Patch-seq experiments, nuclear and cytosolic RNA collected into lysis buffer was reverse transcribed and amplified using SMART-Seq v4 chemistry. Reverse transcription and full-length cDNA amplification were performed with the SMART-Seq v4 Ultra Low Input RNA Kit for Sequencing (Takara, 634894) according to the manufacturer’s instructions, using 21 PCR cycles in 0.65 mL tubes. Reactions were processed in sets of 88 samples. Each amplification set included at least one 8-strip control set containing four no-cell wells and four wells with 10 pg of control RNA. Control RNA consisted of either Mouse Whole Brain Total RNA (Zyagen, MR-201) or control RNA provided with the SMART-Seq v4 kit.

Amplified cDNA was quantified with the Quant-iT PicoGreen dsDNA Assay (Thermo Fisher); yield is reported as total pg in the 12 µL sample. cDNA size distribution was assessed on a 5300 Fragment Analyzer (Agilent) using the HS NGS Fragment Kit (1–6000 bp; Agilent DNF-474) and analyzed with ProSize 4.0 (Agilent).

Amplified cDNA was carried forward into Nextera XT DNA Library Preparation (Illumina FC-131-1096) using either Nextera XT Index Kit v2 Sets A-D (Illumina FC-131-2001, FC-131-2002, FC-131-2003, FC-131-2004) or custom dual indexes from Integrated DNA Technologies. Nextera XT library preparation was performed according to the manufacturer’s instructions except that all reagent volumes, including cDNA input, were reduced to either 0.4× or 0.2× of the standard reaction volume.

### Mapping

Sequence alignment was performed using STAR v2.5.3. Exonic and intronic reads were added together to calculate gene counts, data were analyzed as counts per million reads (CPM). Cells were assigned transcriptomic types using the Hierarchical mapping approach using MapMyCells and is generalized for multiple taxonomies^22,23^. Patch-seq samples were iteratively mapped to nodes: class, subclass and finally supertype/cluster. This iterative approach is repeated 100 times to determine the best fit for each node. Mapping quality was labeled Core, I1, I2, I3 or PoorQ following previously described methods^19,22^; Core and I1 were considered high quality.

### Morphological processing

During Patch-seq experiments, neurons were biocytin-filled via the recording pipette. Slices were fixed in 4% paraformaldehyde in 1× PBS for 1-2 days and stored in PBS at 4 °C. Biocytin-filled neurons were visualized using horseradish peroxidase with DAB chromogen and DAPI staining. Before development, slices were incubated in 1% hydrogen peroxide for 30 min, permeabilized in 2% Triton X-100, and incubated in ABS with 0.1% Triton at 4 °C for 1-2 days, then rinsed.

Imaging followed Allen Institute Patch-seq protocols. Slice overviews were acquired on an AxioImager Z2 (Zeiss) with 20× objective or a SLIDEVIEW VS200 (Evident) with 10× objective. AxioImager overviews used single-plane DAPI and brightfield imaging; VS200 overviews used DAPI and brightfield z-stacks (36-64 μm range, 8-12 μm spacing) collapsed to minimum-intensity projections. High-resolution tiled brightfield z-stacks of identified cells were acquired on the AxioImager Z2 using 63× objectives (1.2 Imm Corr or 1.4 Oil) with 0.44 μm or 0.28 μm z-steps. Stitched 63× image stacks were exported as TIFFs for reconstruction.

All cells were assessed in 20x for the presence of a strong biocytin fill with well-stained dendrites, before proceeding to 63x volume imaging. This assessment stage also served as the measurement of successful morphological recovery throughout this study.

### Standard seal and break-in procedures

Gigaohm sealing and whole cell break-in for standard experiments were performed under MIES control. After contacting the target cell with the pipette, experimenters initiated the seal procedure, which initially switched pressure to −1.4 kPa. Every 20 s, if the steady-state resistance (Rss) slope (least-squares fit over the last 28 test pulses) was below 4 MΩ/s, suction was increased by 0.7 kPa, up to −9.7 kPa, where it was held. Holding potential was set to −70 mV once Rss exceeded 100 MΩ. When Rss reached 1 GΩ, the pipette was vented to atmosphere and the seal was logged. There was no timeout; sealing continued until a gigaseal formed or the operator stopped it.

After the experimenter initiated break-in, suction pulses were delivered from the regulator, with the pipette at atmospheric pressure between them. Each pulse held the regulator at target pressure for 350 ms while opening the valve from regulator to the pipette for the final ∼250 ms. Pulses were at least 5 s apart with 6 different vacuum pressures in sequence: 13.8, 15.5, 20.7, 29.3, 41.4, and 56.9 kPa; then the sequence restarted at the beginning. Break-in was declared when steady-state resistance fell to 1 GΩ or below. There was no pulse limit or timeout.

### Enhanced seal and break-in procedures

Enhanced procesures for establishing gigaohm seal and whole cell break-in were performed under ACQ4 control. The sealing procedure uses an adaptive pressure selection, following observations that seal formation is not strictly improved by applying more pressure; in may cases an optimal intermediate pressure allows the most effective seal formation. The break-in procedure extends the standard method with a more aggressive series of pressure pulses and a more rapid trigger for detecting break-in and terminating pressure pulses.

After contacting the target cell with the pipette, experimenters initiated the seal procedure, which switched to atmospheric pressure for 1 second, followed by an initial suction of −3 kPa, beginning the primary seal-formation control loop. Each iteration of the loop consisted of three steps executed while seal resistance was continuously monitored for the gigaohm success criterion: (1) The applied pressure was swept across a local pressure neighborhood (±2 kPa) in 5-second ramps, both the forward and reverse directions. (2) During the ramps, test pulses were acquired continuously and the maximum change in log(Rss) between consecutive pulses, where each measurement is a 3-point moving average, was identified for each sweep direction independently. The optimal pressure was determined as the mean of these two maxima. (3) Pressure was incremented by 25% of the difference between the current pressure and the measured optimal pressure. This updated pressure was held for 5 seconds before beginning the next iteration.

Once Rss reached at least 100 MΩ (averaged with a 100 ms τ), the holding potential was stepped from 0 mV to −70 mV to minimize cellular stress as the electrode was incorporated into the cellular circuit. If Rss (averaged with a 1 second τ) exceeded 1 GΩ, the seal was considered successful. At this point a pressure of −1 kPa was applied for 5 seconds to allow the seal to fully stabilize before switching to atmospheric pressure. The attempt was aborted if the gigaohm criterion was not met within 80 seconds, or if Rss (averaged with a 10 second τ) stayed below 100 MΩ while increasing less than 1 MΩ/s.

To achieve break-in, a series of negative-pressure pulses was applied, starting with a single −5 kPa, 200 ms pulse and increasing incrementally over 13 rounds to a group of 5 rapid pulses at −60 kPa, 1.5 s each. Rounds were separated by a 2 s waiting period and pulses within a round were separated by a 100 ms gap. Test pulses were acquired continuously and break-in was considered successful when Rss fell below 650 MΩ, at which point the pressure was immediately set to atmospheric. Break-in failed if all 13 rounds passed without a sufficient decrease in Rss, or if the holding current dropped below −1 nA.

### Automated reseal and nucleus extraction

After all electrophysiological recordings were complete, the nucleus was first captured in the pipette tip, followed by extraction by slowly retracting the pipette, allowing the cell membrane to reseal around it. Seal integrity was monitored continuously throughout the procedure via test pulses. The nucleus was then pulled partially or completely inside the pipette to protect it during delivery to its collection tube.

To facilitate initial nuclear capture, the pipette tip was displaced laterally in a series of random steps within the plane perpendicular to the pipette axis, remaining within 2 µm of the starting position, for two consecutive 30 s periods. During each period, suction was ramped linearly from 0 to −2 kPa to begin drawing the nucleus into the pipette tip. Suction was then stepped to −7 kPa, and the pipette was retracted at 0.2 µm/s under voltage clamp at −55 mV.

Membrane integrity during retraction was assessed using two exponential moving averages of resistance with time constants of 5 s and 10 s. The instantaneous rate of change of each average was computed as the log₁₀ ratio of successive averaged values. If either average indicated membrane stretching (rapid increase > 0.01), retraction was paused until the both differences drop below threshold again. If either indicated membrane tearing (any significant decrease), retraction was halted and the pipette was re-advanced until resistance recovered. If the slow average fell below 50% of the pre-retraction baseline, indicating irreversible disruption, the procedure was aborted.

Reseal was considered successful when either the total retraction distance exceeded 200 µm, or the pipette had retracted at least 20 µm and resistance had exceeded four times the pre-retraction baseline for at least 5 consecutive seconds. Upon success, the pipette was withdrawn from the tissue at 6 µm/s, suction was increased to −10 kPa for 30 s (stopping early if Rss drops below 20 MΩ) to draw the nucleus fully into the pipette lumen, and the imaging focal plane was set to the pipette tip for visual confirmation. If reseal criteria were not met within 10 minutes, the procedure was aborted.

### Automated nucleus collection

Nuclear material, once attached to the pipette tip and exposed to air, becomes tough and firmly adheres to the pipette. We used sonication to expel this material into a collection tube without breaking the pipette tip. Sonication was produced using a custom-fabricated actuator assembly (Fig. 6A) incorporating a piezoelectric chip (Thorlabs, PA3CKW) that is similar to one previously developed for pipette cleaning^20^. The actuator was driven by an analog output from a data acquisition (DAQ) device (NI USB-6351), which was then amplified by a fixed-gain voltage amplifier (PiezoDrive, PD200). The cap housing the piezoelectric chip was designed to interface with a standard electrode holder (Molecular Devices, HL-U) and was fabricated using stereolithographic 3D printing with clear resin (Formlabs, Form 4). A flexible internal housing and washer for the piezoelectric chip were fabricated using Flexible 80A resin (Formlabs, Form 4) to ensure stable mechanical coupling and alignment. Epoxy (Henkel, Loctite M-121HP) was applied between the piezoelectric chip and the cap, and between the cap and the cap adapter, to secure the components in place. Once the nucleus was captured, it was deposited into a PCR tube containing buffer solution at a dedicated PCR tube station (Fig. 6B).

**Figure 6.**
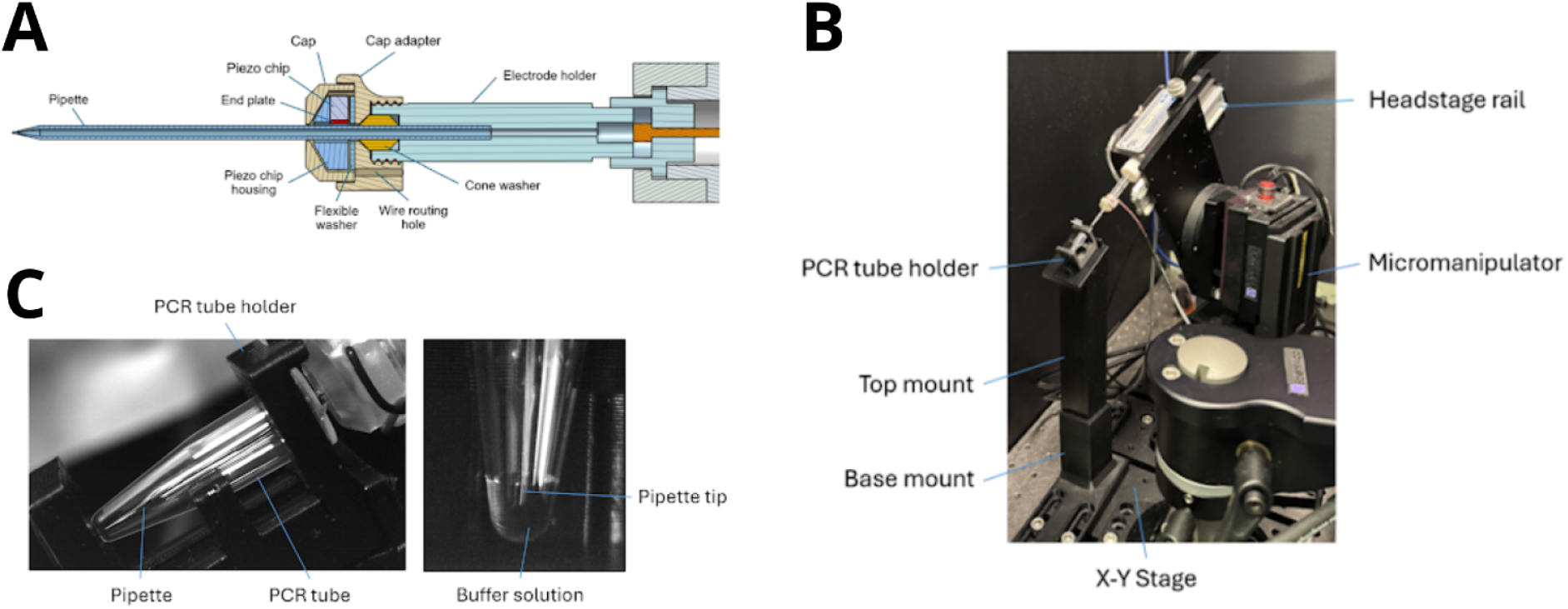
Nucleus collection hardware. **A.** CAD model of the electrode holder and integrated sonicator cap assembly (Onshape). **B.** Nucleus is deposited into a PCR tube at a dedicated collection station. **C.** Pipette positioned at the pre-calibrated nucleus collection location within the PCR tube (left). Top view of the pipette tip immersed in buffer solution (right).

To enable repeatable positioning during this process, a custom PCR tube holder and mounting system were designed and 3D-printed using PETG-HF (Bambu Lab, H2D). The mount consists of a base fixed to the X-Y stage (Scientifica, MMBP), maintaining a constant spatial relationship with the micromanipulator holding the headstage and pipette. A removable top mount is magnetically coupled to the base to allow rapid installation and removal.

The top mount incorporates a rail-guided PCR tube holder that slides into position and locks via a magnetic stop at a 29° angle, matching the pipette approach angle to facilitate efficient nucleus transfer. The system is positioned such that the pipette tip can reliably access and contact the buffer solution within the PCR tube (Fig. 6C). During collection, the nucleus was retained at the pipette tip using a holding pressure of −2 kPa and returned to a home position. The headstage was then raised along its rail (Scientifica) and secured at the upper position, after which the micromanipulator rotary base (Scientifica) was manually rotated to a fixed magnetic stop to align the pipette with the PCR tube station. The pipette was then automatically driven to a pre-calibrated location within the tube, ensuring the tip was fully immersed in the buffer solution. Due to the small quantity of buffer solution (∼10 μl), reliably hitting this small target requires a per-pipette calibration of the tip’s position relative to the manipulator’s coordinate system.

A simultaneous pressure / sonication sequence was then applied to release the nucleus: 68 kPa for 5 s and −35 kPa for 2 s, iterated five times to agitate the nucleus and introduce lysis buffer into the pipette, followed by 68 kPa for 130.0 s to eject most of the pipette contents. Sonication was applied for a total of 160 s with eight consecutive frequency-sweep cycles. Each sonication cycle consisted of a 10 s upsweep from 90 to 155 kHz, followed by a 10 s downsweep from 155 to 90 kHz. The signal was maintained at 36 Vpp with an 18 V DC offset. The sonication frequency was tuned near resonance to enable rapid nucleus release while minimizing disruption to nucleus content.

### Pipette cleaning

Pipette cleaning was performed by pulsing high pressure through the pipette while immersed in an enzymatic detergent solution^16^ (Tergazyme, Alconox; Fig. 7). The sonicator used for nucleus collection simultaneously provided mechanical agitation of the pipette to remove detergent-resistant nuclear material. Detergent was gravity-fed from a reservior to a small well that was surrounded by an aspirated moat to prevent cross contamination of detergent into the recording chamber. Detailed assembly and system design are described in our protocol^24^.

**Figure 7.**
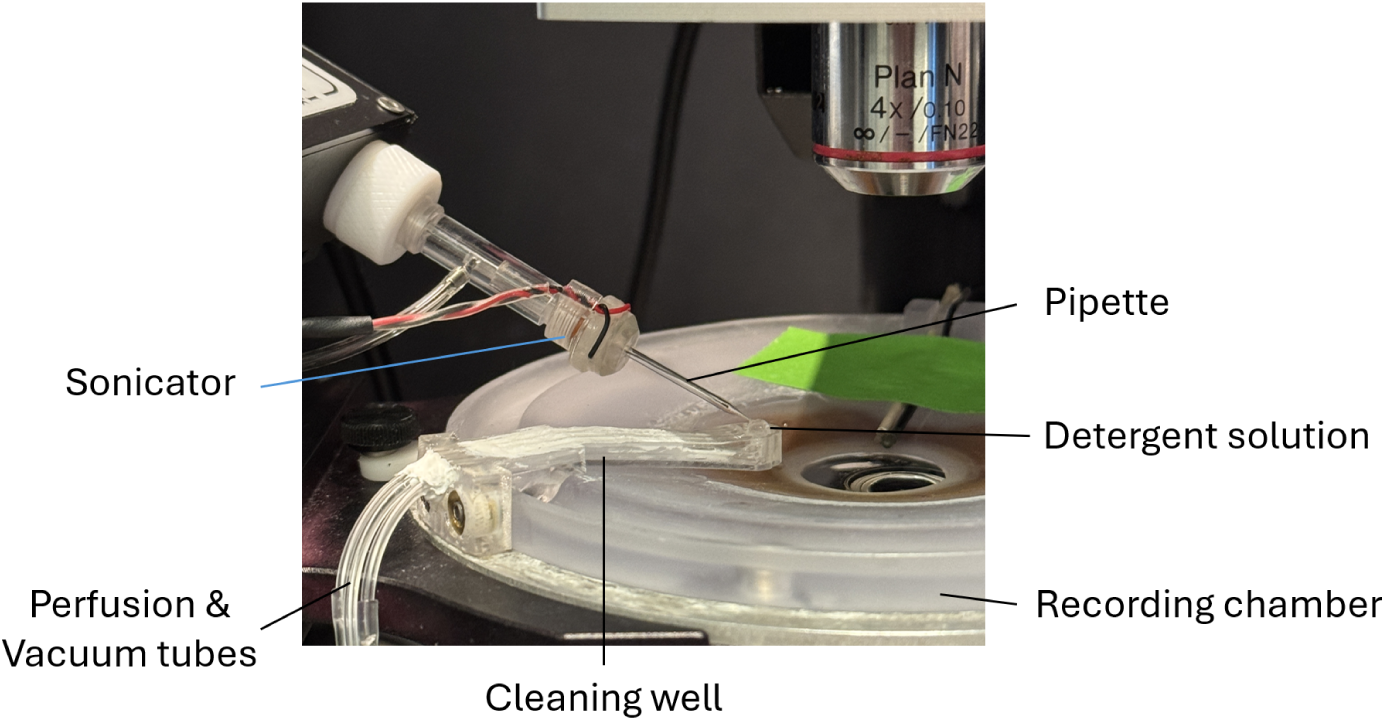
Automated pipette cleaning. The pipette tip, secured in the electrode holder capped with a sonicator, is immersed in the detergent solution in the cleaning well.

Following unsuccessful patch attempts or nucleus extraction, the pipette was repositioned to a pre-calibrated location in the cleaning well, ensuring the tip was fully immersed. Cleaning was achieved through a combination of controlled pressure cycling and sonication to remove debris and residual biological material from both the interior and exterior surfaces of the pipette.

Pressure pulses were delivered using a pressure regulation system (Proportion Air, QPV1), consisting of alternating negative and positive pressures as follows: 68 kPa for 3 s and −30 kPa for 2.0 s, iterated five times, followed by 68 kPa for 30.0 s.

Simultaneously, sonication was applied using a bidirectional frequency sweep near the resonance of the coupled piezo–pipette system to rapidly remove contaminants from the pipette. The sweep consisted of a 23 s upsweep from 90 to 155 kHz, followed by a 23 s downsweep from 155 to 90 kHz. The signal was maintained at 36 V peak-to-peak (Vpp) with an 18 V DC offset.

During experiments, the cleaning solution level was maintained via pressure equilibration with an external reservoir, ensuring a constant meniscus within the well. A convex meniscus was used to improve visualization and assist pipette tip positioning during immersion into the cleaning solution. It also minimized the risk of collision with the cleaning well wall under tight spatial constraints. Excess solution was continuously removed through a vacuum line to prevent overflow into the recording chamber.

Following completion of the cleaning cycle, the pipette was withdrawn from the well and repositioned at the recording site for subsequent use.

